# TaxoFormer: Hierarchical Transformer for Predicting the Full Taxonomic Lineage of Protein Sequences

**DOI:** 10.64898/2026.06.06.730618

**Authors:** Mohammad Parsa, Kooshiar Azimian, Kathy Y. Wei

## Abstract

Predicting labels in massive, hierarchically structured output spaces is a core challenge in machine learning. In this work, we use the problem of predicting the full taxonomic lineage of a protein from its sequence as a case study for this challenge. We introduce TaxoFormer, an architecture whose primary contribution is a structured tokenization scheme that losslessly represents the entire NCBI phylogenetic tree, a graph with over 1.3 million nodes using a compact vocabulary of just 15,000 tokens. By coupling a pre-trained ESM-2 model with an autoregressive decoder and training with a standard cross-entropy objective, we test the hypothesis that a simple generative objective is sufficient to learn complex, latent structure when the output space is explicitly modeled. We show that this approach is highly effective: on a dataset of 188 million proteins, the model not only achieves accurate lineage prediction but also implicitly learns a continuous, phylogenetically-structured latent space. This work provides a scalable, alignment-free method for taxonomic annotation and demonstrates that explicitly modeling the structure of a complex output space is a powerful mechanism for learning meaningful representations.^2^

## 1 Introduction

Many real-world prediction problems, from product categorization to medical diagnosis and scientific discovery, involve labels that are not independent but exist in a massive, hierarchically structured space. Training models to navigate these spaces is a fundamental challenge for machine learning. A naive classification approach with a flat softmax is computationally intractable and ignores the rich relational information embedded in the hierarchy.

In this work, we explore this challenge through the lens of computational biology. The taxonomic “tree of life” is a canonical example of such a space, with over 1.3 million nodes (taxa) organized into a deep, variable-depth hierarchy. Assigning a protein sequence to its correct position in this tree is a critical task, as a protein’s lineage is a powerful prior for its function and structure. While traditional methods exist, they often rely on computationally intensive multiple sequence alignments (MSAs) [4, 15] and tree-building algorithms [11, 9], and fail on novel sequences without close homologs.

We formulate this biological problem as a general machine learning task: generating a variable-length path through a massive, hierarchical graph from a single input sequence. To address this, we propose TaxoFormer, an encoder-decoder framework whose core innovation is a hierarchical tokenization scheme that provides a strong inductive bias by explicitly modeling the structure of the output space. We demonstrate that this architectural choice, when combined with a simple cross-entropy objective, is sufficient to induce the learning of a powerful, phylogenetically-aware representation space as an emergent property.

## 2 Related Work

Our work builds upon large-scale PLMs like ESM-2 [6], which learn rich protein representations but are not explicitly designed to generate labels within a structured space. The challenge of representing a massive, structured label space is analogous to open-vocabulary problems in NLP [13]. While some methods learn hierarchical representations from unstructured text [7, 16], we leverage an explicit, known symbolic hierarchy—the phylogenetic tree—to structure the output token space. A recent line of work distinguishes between “evolutionary modeling” (e.g., fixed-rank prediction) and “evolutionary reasoning”—the ability to infer complex, hierarchical relationships [14]. While deep learning has been applied to phylogenetics with methods like VAEs [19] and autoregressive models [20], these often operate on existing alignments or trees. In contrast to models that reconstruct a tree from a *set* of unaligned sequences [14, 8], TaxoFormer approaches this from a different perspective: generating the full path for a *single* sequence using a standard cross-entropy objective.

## 3 Methodology

We formulate lineage prediction as a sequence-to-sequence task, illustrated in Figure 1. The core of our approach is a method to encode the *≈*1.39 million taxonomic nodes into a compact, fixed-length sequence of 118 tokens. This is processed by an encoder-decoder model, which uses a pre-trained ESM-2 model as its encoder and a 4-layer Transformer as its decoder, adapted with QLoRA for parameter-efficient fine-tuning. Full technical details are provided in Appendix A.

**Figure 1:**
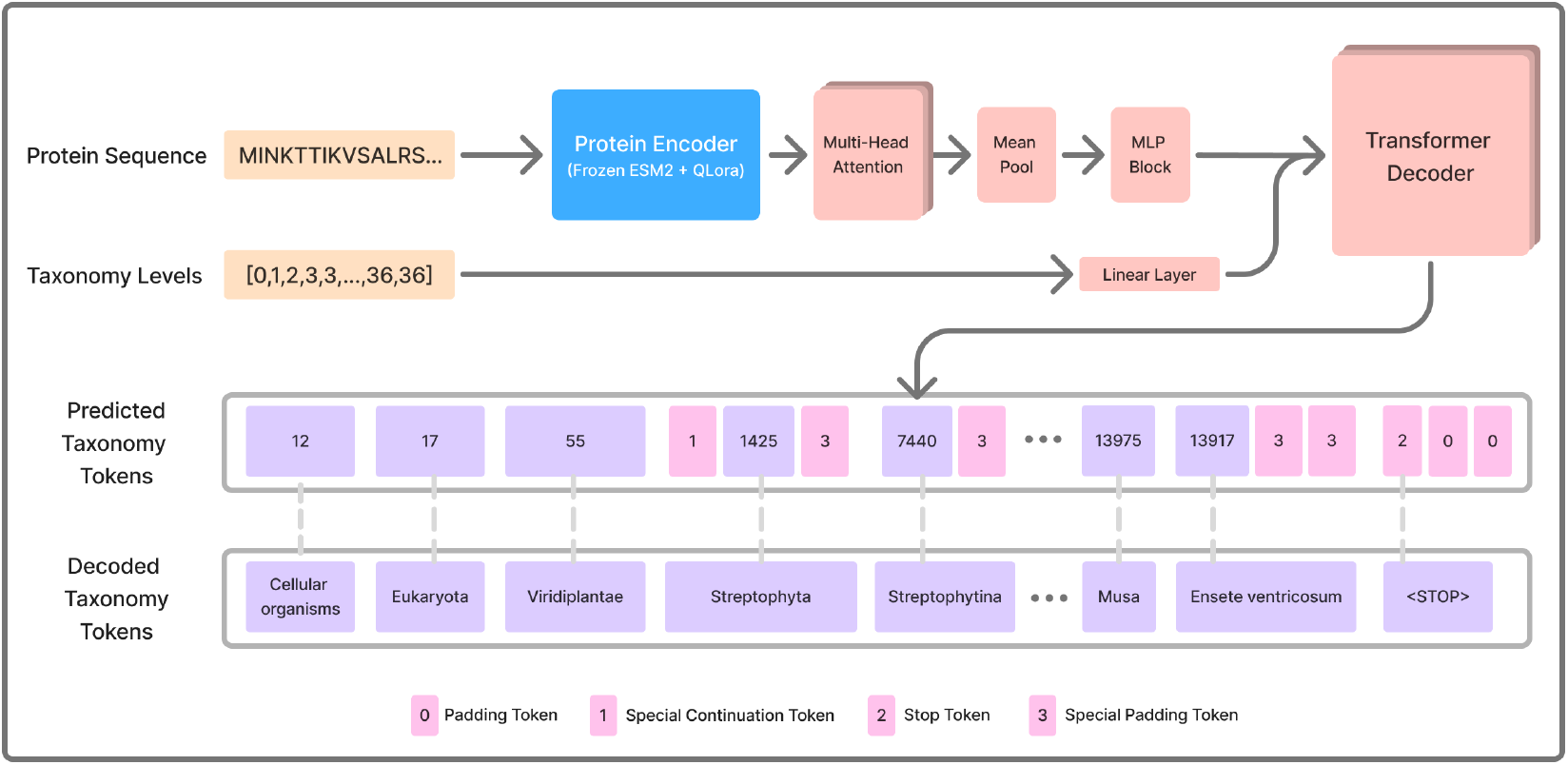
TaxoFormer Architecture. A protein sequence is fed into the frozen ESM-2 encoder with trainable QLoRA adapters. A multi-head attention pooling layer distills the sequence into a single embedding, which serves as cross-attention context for the autoregressive Transformer decoder. The decoder generates a sequence of 118 integer tokens, which are then converted via deterministic logic into the final, human-readable taxonomic lineage.

## 4 Experiments and Results

### 4.1 Dataset and Implementation

The scale and structure of our biological data, shown in Figure 2, provide a challenging testbed for learning on hierarchical label spaces. We constructed our dataset from UniProt [17] (*≈* 188 million proteins after filtering), curating a 100,000-record evaluation subset using a deterministic, stratified systematic sampling scheme. A crucial point is that the results shown are from a checkpoint after training on only 34% of the total training data. The strong performance reported here, achieved with the model in a partially trained state, demonstrates the sample efficiency of our approach and suggests significant potential for improvement as training concludes.

**Figure 2:**
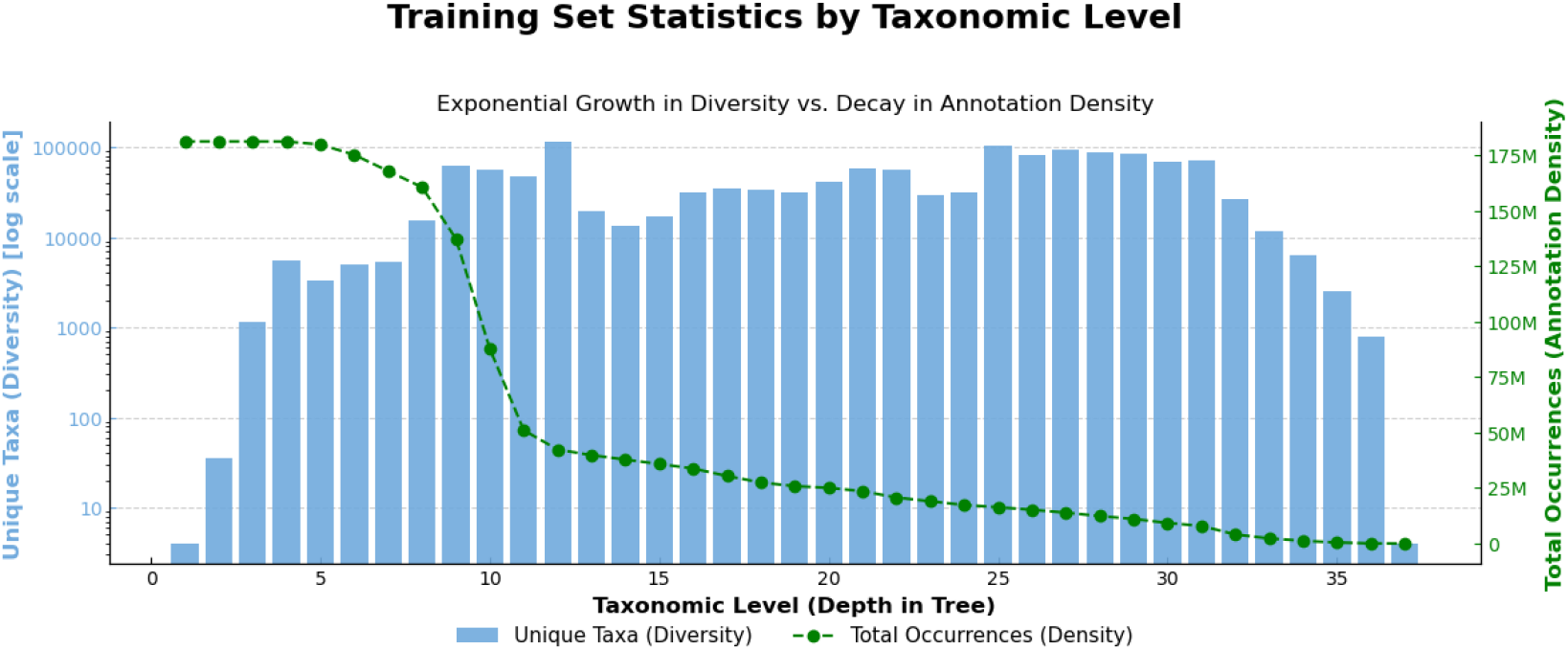
The Hierarchical Label Space Challenge. This visualization of the training set illustrates the core problem. The number of unique taxa (blue bars, log scale) grows exponentially at deeper levels, creating a vast label space. Simultaneously, the number of sequences with annotations at those depths (green line) decays, leading to data sparsity. The model must handle both exploding diversity and decreasing data density.

### 4.2 Quantitative Results and Generalization

Figure 3 provides a comprehensive overview of model performance. The model demonstrates robust generalization to novelty; Figure 3D shows the performance drop on sequences containing taxa never seen during training is minor. This is a key strength for real-world scientific applications. Figure 3C validates our core hypothesis: the strong positive correlation between token accuracy (our direct training objective) and the inverted RF distance (a measure of phylogenetic correctness) confirms that optimizing a simple generative objective on a well-structured output space directly leads to learning the complex, latent structure of the problem.

**Figure 3:**
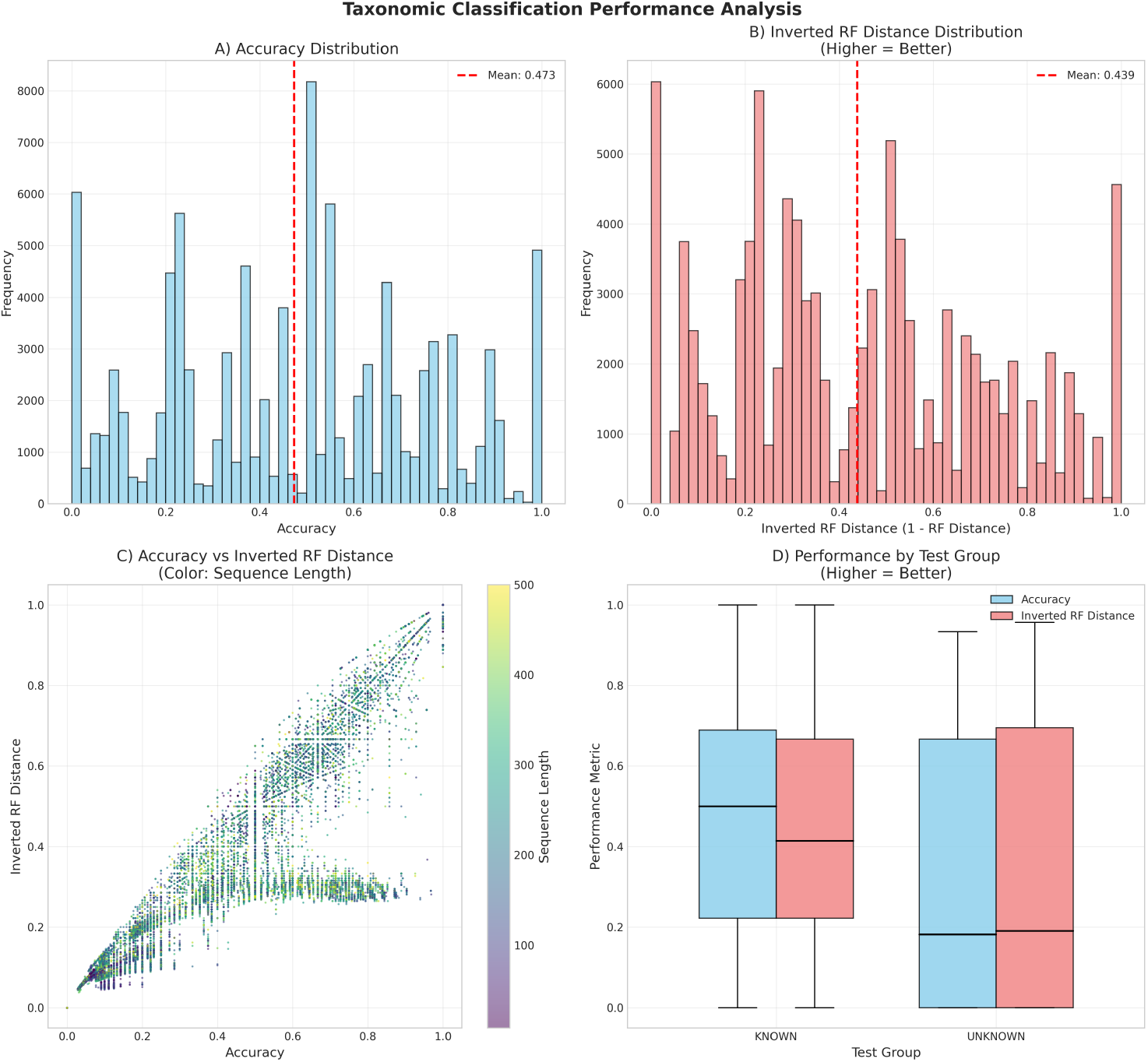
Overall Performance Analysis. (A) Distribution of token-level accuracy. (B) Distribution of inverted Robinson-Foulds (RF) distance (1 - RF). (C) A strong correlation between accuracy and inverted RF distance. (D) A comparison of performance on sequences with only known vs. at least one novel taxon, demonstrating robust generalization.

### 4.3 Implicit Learning of Phylogenetic Structure on Benchmarks

As shown in Table 1, TaxoFormer performs competitively on multi-sequence reconstruction benchmarks against PHYLA [14], a specialized model. Our result is noteworthy because TaxoFormer was trained on a different task with a simpler objective, yet implicitly learns sufficient structure to be competitive under a more challenging end-to-end evaluation protocol, even surpassing PHYLA on the TreeBase benchmark.

**Table 1:** Benchmark Performance Comparison. Normalized Robinson-Foulds (RF) distance on TreeFam and TreeBase (lower is better, ↓).

| Model | Parameters | Evaluation Method | TreeFam ↓ | TreeBase ↓ |
| --- | --- | --- | --- | --- |
| Hamming Distance | – | Embed → NJ* | 0.7509 | 0.7466 |
| ESM-2 [6] | 650M | Embed → NJ* | 0.6669 | 0.7809 |
| ProGen2-XLarge [10] | 6.4B | Embed → NJ* | 0.8249 | 0.8624 |
| Evo 2 [1] | 7B | Embed → NJ* | 0.8406 | 0.8372 |
| PHYLA [14] | 24M | Embed → NJ* | **0.5777** | 0.7269 |
| **TaxoFormer (Ours)** | 802M | **Direct Lineage Generation** | 0.7010 | **0.7240** |

\*Tree constructed from sequence embeddings via the Neighbor-Joining (NJ) algorithm.

### 4.4 Analysis of the Learned Evolutionary Landscape

To probe the structure of the learned representation space, we visualized synthetic evolutionary trajectories (Figure 4). We selected orthologous protein pairs (source organism to human) and generated intermediate sequences by iteratively mutating the source toward the target. Each intermediate sequence was passed through TaxoFormer’s frozen encoder to obtain its embedding, and all resulting embeddings from these paths were projected onto a 2D manifold using t-SNE.

**Figure 4:**
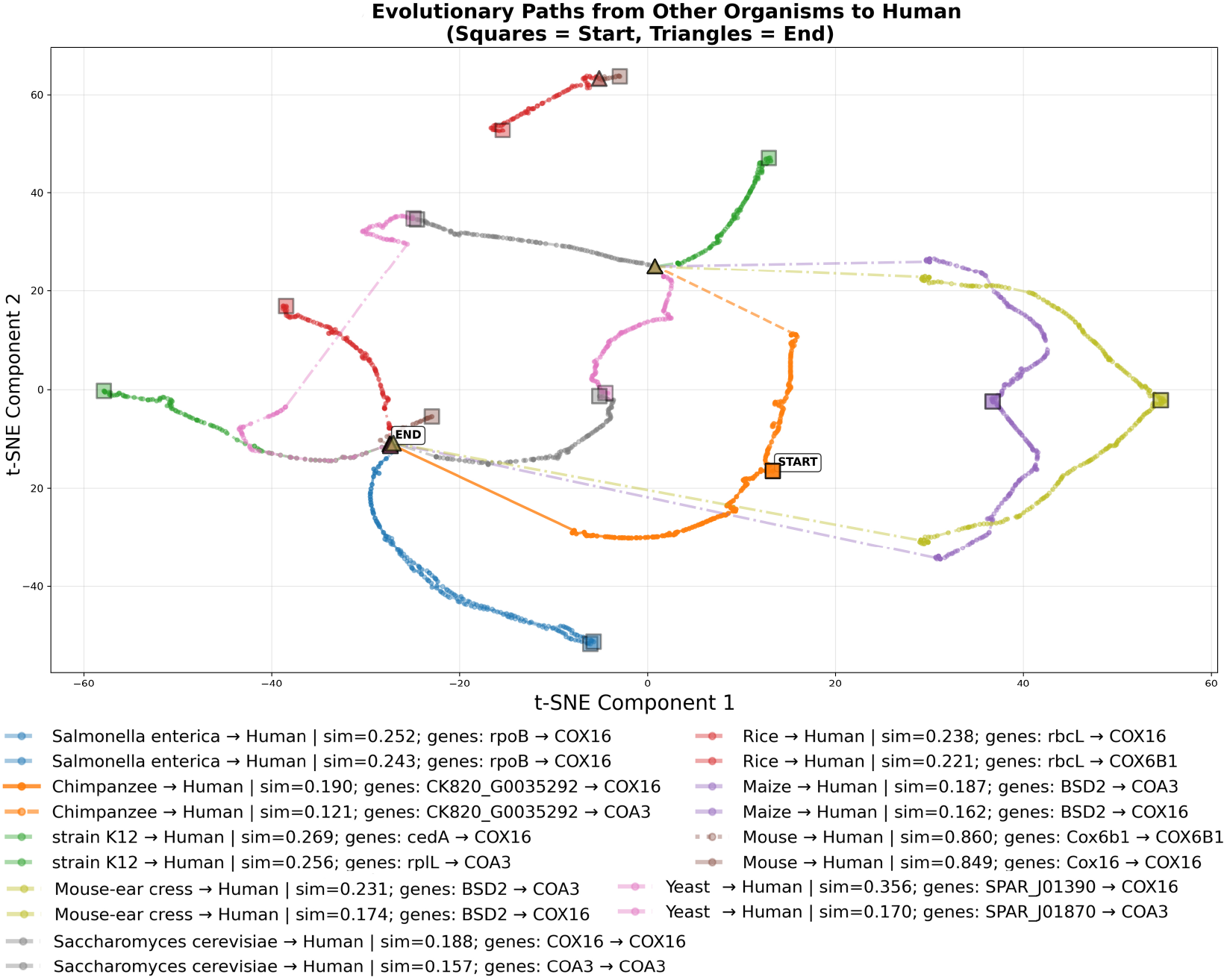
Learned Evolutionary Trajectories. A t-SNE visualization of protein embeddings along synthetic evolutionary paths to *Homo sapiens*. The smooth, directed trajectories indicate a well-structured, evolution-aware latent space. While t-SNE is excellent for visualizing local structure, inter-cluster distances are not necessarily meaningful.

The visualization reveals that the paths are not random walks, but rather smooth, coherent trajectories that progress directionally. The observed smoothness is critical: it demonstrates a well-organized latent space where single mutations map to contiguous steps in the embedding. This directed nature suggests the model has learned a continuous “phylogenetic gradient,” where proximity and direction correspond to evolutionary relatedness. This is a powerful emergent property, as this continuous manifold was learned solely from the task of predicting discrete taxonomic labels, highlighting the power of our overall approach.

## 5 Discussion

Our results present a cohesive picture: the quantitative metrics (Figure 3) and competitive benchmarks (Table 1) demonstrate *what* the model can do, while the latent space analysis (Figure 4) reveals *how* it does so—by implicitly learning a structured evolutionary landscape. Our central finding is that a simple, generative cross-entropy objective, when paired with a strong inductive bias from a hierarchical tokenizer, is a remarkably effective signal for learning a phylogenetically meaningful latent space.

The model’s errors are not random noise but are themselves informative. As shown in Figure 5, performance is correlated with data availability. More interestingly, a qualitative analysis reveals that many errors are biologically plausible. For instance, the model frequently confuses sister genera within the same family, indicating it has learned fine-grained relationships. In other cases, a viral protein may be misclassified into its host’s lineage, a potential signature of viral mimicry or genomic integration, a phenomenon observed in archaeal genomes as well [5]. This suggests the model’s “errors” could be used as a tool for discovering interesting evolutionary events.

**Figure 5:**
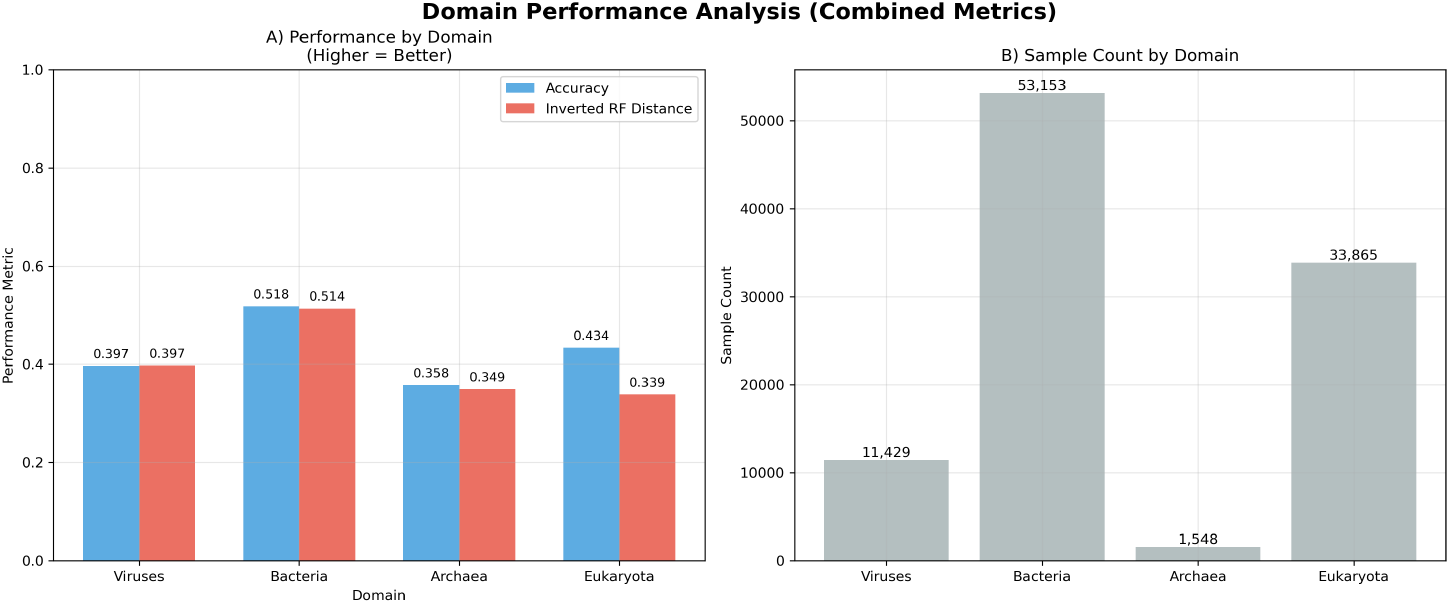
Performance Breakdown by Domain on the Evaluation Subset. This figure illustrates the model’s performance across the major domains of life within the evaluation set. Our stratified sampling method was designed to ensure this subset mimics the class distribution of the full training set. Therefore, the heavy imbalance shown in subplot (B), with Bacteria and Eukaryota vastly outnumbering Viruses and Archaea, faithfully reflects the data landscape the model was trained on. Subplot (A) shows that performance is strongly correlated with sample count. Notably, despite Archaea having the smallest representation in the evaluation set, the model maintains a respectable predictive accuracy of 36%, demonstrating robust generalization on underrepresented classes.

This study is based on a single training run; future work should include multiple seeds to establish statistical confidence. A key next step is to conduct ablation studies on core design choices. Furthermore, the model does not offer calibrated uncertainty estimates, a critical feature for deployment.

## 6 Conclusion

In this work, we used the problem of protein taxonomy to explore the broader machine learning challenge of learning on massive, hierarchically structured label spaces. We demonstrated that by explicitly modeling the structure of the output space with a hierarchical tokenizer, a standard encoder-decoder model trained with a simple objective can implicitly learn the complex, latent grammar of the domain. Our model, TaxoFormer, achieves competitive performance and robust generalization, highlighting this as a promising and elegant strategy for tackling complex scientific and real-world prediction problems.

## A Detailed Methodology

### A.1 Hierarchical Taxonomic Encoding

The central innovation of our work is a method to encode the vast and sparse NCBI taxonomic tree, with its ≈ 1.39 million nodes across 37 levels, into a dense, fixed-length format. Instead of a flat vocabulary, we create a structured one. We represent the phylogenetic tree with 37 distinct levels (e.g., Kingdom, Phylum, Class) and map these to a sequence of 118 “bins.” Each bin is a distinct vocabulary space of size 15,000. If a taxonomic level contains fewer than 15,000 unique nodes (taxa), it is mapped to a single bin. If it contains more (e.g., the “Genus” level), its nodes are distributed alphabetically across multiple consecutive bins. A positional encoding is maintained so the model knows which bin(s) correspond to each taxonomic level.

The final encoded sequence for a lineage is a list of 118 integers, generated by placing a taxon’s corresponding token ID into its assigned bin. We use three special tokens:

- [`PAD`] (0): Used for all unused positions in the 118-token sequence.
- [`CONT`] (1): A continuation token. If a level spans multiple bins, this token links them, signaling the decoder to continue generating tokens for the same taxonomic level.
- [`STOP`] (2): Marks the end of a taxonomic path, allowing for variable-depth lineages and indicating no further known classification.

This binned representation drastically reduces the vocabulary size from over a million to just 15,000, making the generative task computationally feasible while retaining the capacity to represent the entire tree losslessly.

### A.2 Model Architecture

TaxoFormer uses an encoder-decoder architecture totaling 802M parameters.

#### Protein Encoder

The encoder is the pre-trained ESM-2 model from the 650M parameter class [6]. To efficiently adapt this large model, we freeze its base parameters and apply QLoRA [2], a parameter-efficient fine-tuning technique. We insert Low-Rank Adaptation (LoRA) [3] adapters with a rank of 8 to the query, key, and value matrices of the self-attention layers. These adapters constitute only *≈*8M of the encoder’s 659M parameters.

#### Pooling Layer

To distill the variable-length sequence representation from the encoder into a single fixed-size vector suitable for the decoder, we pass the final-layer embeddings through a multi-head attention pooling layer. We use the [CLS] token embedding as the query to the pooling layer, which attends to all other token embeddings in the sequence to produce a final, context-aware protein embedding **z** ∈ ℝ^*d*^.

#### Taxonomy Decoder

The decoder is a standard 4-layer Transformer architecture [18] with 143M parameters. It takes the protein embedding **z** as cross-attention context and autoregressively generates the 118-token encoded taxonomy sequence *Y* = (*y*_1_, …, *y*_118_). Specifically, the embedding **z** is provided as the key and value to each of the decoder’s cross-attention layers, allowing the decoder at each step to focus on the most relevant aspects of the protein’s representation when predicting the next taxonomic token. In total, only 151M of the 802M parameters (the decoder and the LoRA adapters) are trainable.

### A.3 Training Objective

We deliberately employ a standard cross-entropy loss (ℒ_CE_). We hypothesize that a standard generative objective is a sufficient supervisory signal to force the model to learn the complex, hierarchical grammar of the tree of life as an implicit property, without needing specialized, tree-aware loss functions. The model is trained end-to-end by minimizing ℒ_CE_ between the predicted token probabilities and the ground-truth encoded taxonomy sequence, ignoring the `[PAD]` token index.

## B Supplementary Context

### B.1 Detailed Related Work

Our work builds upon large-scale PLMs like ESM-1b [12], ESM-2 [6], and ProGen2 [10], which learn rich protein representations through self-supervision on massive sequence corpora. While these models excel as feature extractors for downstream tasks, they are not explicitly designed to navigate or generate labels within a structured, hierarchical space like the complete tree of life.

The challenge of representing a massive, structured label space is analogous to open-vocabulary problems in NLP. Standard subword tokenizers [13] create large, static vocabularies that are inflexible to new domains. Recent work has explored hierarchical or “tokenizer-free” models that build representations dynamically [7, 16]. While these methods learn hierarchical representations from unstructured text, TaxoFormer introduces a complementary approach: we leverage an explicit, known symbolic hierarchy—the phylogenetic tree—to structure the output token space for a generative model.

### B.2 Dataset Curation Details

To create our validation set, we sample a deterministic, systematic subset of the full 7-million-record test split. This procedure is seedless and fully reproducible. The universe for sampling, *U*, consists of all records in the test split. We define seven depth buckets, *B*_1_, …, *B*_7_, and form strata *S*_*G,B*_ as the Cartesian product of the top-level lineage *G* ∈ *{*Viruses, Cellular organisms*}* and the depth bucket *B*. For a target subset size of 100,000 records, each stratum receives a proportional allocation. Within each stratum, we impose a fixed lexicographic order based on the full lineage string and select records at evenly spaced intervals using systematic sampling.

## C Prediction Decoding Logic and Examples

The following five examples are drawn from our 100k evaluation subset and illustrate the decoding process in practice.

### C.1 Example 1

- **Ground Truth Tree:** Cellular organisms, Eukaryota, Viridiplantae, Streptophyta, Strep-tophytina, Embryophyta, Tracheophyta, Euphyllophyta, Spermatophyta, Magnoliopsida, Mesangiospermae, Liliopsida, Petrosaviidae, commelinids, Zingiberales, Musaceae, Ensete, Ensete ventricosum (Abyssinian banana) (Musa ensete)
- **Raw Ground Truth Tokens:** [12, 17, 55, 1425, 7440, 11765, 2031, 7482, 0, 8074, 0, 0, 0, 0, 13975, 0, 0, 0, 13744, 0, 0, 0, 8013, 0, 0, 0, 0, 0, 0, 0, 0, 0, 14714, 0, 5156, 3795, 0, 6183, 0, 0, 8353, 0, 0, 13917, 0, 0, 2, 0, 0, 0, 0, 0, 0, 0, 0, 0, 0, 0, 0, 0, 0, 0, 0, 0, 0, 0, 0, 0, 0, 0, 0, 0, 0, 0, 0, 0, 0, 0, 0, 0, 0, 0, 0, 0, 0, 0, 0, 0, 0, 0, 0, 0, 0, 0, 0, 0, 0, 0, 0, 0, 0, 0, 0, 0, 0, 0, 0, 0, 0, 0, 0, 0, 0, 0, 0, 0, 0, 0]

- **Raw Predicted Tokens:** [12, 17, 55, 1425, 7440, 11765, 2031, 7482, 8074, 8074, 13975, 13975, 13975, 17, 13975, 17, 13744, 13744, 13744, 8011, 8011, 8011, 8011, 16, 14714, 14714, 14714, 12, 16, 12, 14714, 14714, 14714, 3869, 5156, 3785, 6201, 6183, 8359, 8359, 8359, 8359, 8359, 13917, 2, 2, 2, 2, 2, 2, 2, 2, 2, 2, 2, 2, 2, 2, 2, 2, 2, 2, 2, 2, 2, 2, 2, 2, 2, 2, 2, 2, 2, 2, 2, 2, 2, 2, 2, 2, 2, 2, 2, 2, 2, 2, 2, 2, 2, 2, 2, 2, 2, 2, 2, 2, 2, 2, 2, 2, 2, 2, 2, 2, 2, 2, 2, 2, 2, 2, 2, 2, 2, 2, 2, 2, 2, 2]
- **Decoded Prediction Tree:** Cellular organisms, Eukaryota, Viridiplantae, Streptophyta, Streptophytina, Embryophyta, Tracheophyta, Euphyllophyta, Spermatophyta, Magnoliopsida, Mesangiospermae, eudicotyledons, Petrosaviidae, commelinids, Poales, Musaceae, Musa, Ensete ventricosum (Abyssinian banana) (Musa ensete)

### C.2 Example 2

- **Ground Truth Tree:** Cellular organisms, Eukaryota, Opisthokonta, Fungi, Dikarya, Basidiomycota, Agaricomycotina, Tremellomycetes, Tremellales (jelly fungi), Cuniculitremaceae, Kockovaella, Kockovaella imperatae
- **Raw Ground Truth Tokens:** [12, 17, 53, 1423, 7437, 11778, 2041, 7564, 0, 8133, 0, 0, 0, 0, 203, 0, 0, 0, 14199, 0, 0, 0, 8402, 0, 0, 0, 0, 0, 0, 0, 0, 0, 2, 0, 0, 0, 0, 0, 0, 0, 0, 0, 0, 0, 0, 0, 0, 0, 0, 0, 0, 0, 0, 0, 0, 0, 0, 0, 0, 0, 0, 0, 0, 0, 0, 0, 0, 0, 0, 0, 0, 0, 0, 0, 0, 0, 0, 0, 0, 0, 0, 0, 0, 0, 0, 0, 0, 0, 0, 0, 0, 0, 0, 0, 0, 0, 0, 0, 0, 0, 0, 0, 0, 0, 0, 0, 0, 0, 0, 0, 0, 0, 0, 0, 0, 0, 0, 0]
- **Raw Predicted Tokens:** [12, 17, 55, 1423, 7437, 11766, 2106, 7564, 8133, 8133, 14005, 14005, 14005, 17, 14005, 17, 14199, 14199, 14199, 14199, 8402, 8402, 8402, 14987, 2, 2, 2, 12, 17, 12, 2, 2, 2, 2, 2, 2, 2, 2, 2, 2, 2, 2, 2, 2, 2, 2, 2, 2, 2, 2, 2, 2, 2, 2, 2, 2, 2, 2, 2, 2, 2, 2, 2, 2, 2, 2, 2, 2, 2, 2, 2, 2, 2, 2, 2, 2, 2, 2, 2, 2, 2, 2, 2, 2, 2, 2, 2, 2, 2, 2, 2, 2, 2, 2, 2, 2, 2, 2, 2, 2, 2, 2, 2, 2, 2, 2, 2, 2, 2, 2, 2, 2, 2, 2, 2, 2, 2, 2]
- **Decoded Prediction Tree:** cellular organisms, Eukaryota, Viridiplantae, Fungi, Dikarya, Ascomycota, Pucciniomycotina, Tremellomycetes, Tremellales (jelly fungi), Cryptococcaceae, Kockovaella, Kockovaella imperatae

### C.3 Example 3

- **Ground Truth Tree:** Cellular organisms, Eukaryota, Viridiplantae, Streptophyta, Strep-tophytina, Embryophyta, Tracheophyta, Euphyllophyta, Spermatophyta, Magnoliopsida, Mesangiospermae, eudicotyledons, Gunneridae, Pentapetalae, asterids, lamiids, Boraginales, Hydrophyllaceae, Nemophila, Nemophila menziesii (Baby blue eyes)
- **Raw Ground Truth Tokens:** [12, 17, 55, 1425, 7440, 11765, 2031, 7482, 0, 8074, 0, 0, 0, 0, 13975, 0, 0, 0, 13744, 0, 0, 0, 8011, 0, 0, 0, 0, 0, 0, 0, 0, 0, 14712, 0, 5153, 3786, 0, 6172, 0, 0, 8744, 0, 0, 14332, 0, 0, 5641, 0, 0, 10191, 0, 0, 2, 0, 0, 0, 0, 0, 0, 0, 0, 0, 0, 0, 0, 0, 0, 0, 0, 0, 0, 0, 0, 0, 0, 0, 0, 0, 0, 0, 0, 0, 0, 0, 0, 0, 0, 0, 0, 0, 0, 0, 0, 0, 0, 0, 0, 0, 0, 0, 0, 0, 0, 0, 0, 0, 0, 0, 0, 0, 0, 0, 0, 0, 0, 0, 0, 0]
- **Raw Predicted Tokens:** [12, 17, 55, 1425, 7440, 11765, 2031, 7482, 8074, 8074, 8074, 13975, 8074, 17, 13975, 17, 13744, 13744, 13744, 8074, 8011, 8011, 8011, 17, 14712, 14712, 14712, 12, 17, 12, 14712, 14712, 14712, 8417, 5153, 3783, 6172, 6172, 8337, 8337, 8337, 8337, 14152, 14152, 2, 2, 14942, 8363, 6791, 6791, 2, 8104, 2, 2, 2, 2, 2, 2, 2, 2, 2, 2, 2, 2, 2, 2, 2, 2, 2, 2, 2, 2, 2, 2, 6892, 2, 2, 2, 2, 2, 2, 2, 2, 2, 2, 2, 2, 2, 2, 2, 2, 2, 2, 2, 2, 2, 2, 2, 2, 2, 2, 2, 2, 2, 2, 2, 2, 2, 2, 2, 2, 2, 2, 2, 2, 2, 2, 2]
- **Decoded Prediction Tree:** Cellular organisms, Eukaryota, Viridiplantae, Streptophyta, Streptophytina, Embryophyta, Tracheophyta, Euphyllophyta, Spermatophyta, Magno-liopsida, Mesangiospermae, eudicotyledons, Gunneridae, Pentapetalae, rosids, lamiids, Solanales, Boraginaceae

### C.4 Example 4

- **Ground Truth Tree:** Cellular organisms, Archaea, Euryarchaeota, Methanomada group, Methanobacteria, Methanobacteriales, Methanobacteriaceae, Methanobrevibacter, Methanobrevibacter millerae
- **Raw Ground Truth Tokens:** [12, 18, 58, 1458, 7521, 11886, 2164, 7706, 0, 13469, 0, 0, 0, 0, 2, 0, 0, 0, 0, 0, 0, 0, 0, 0, 0, 0, 0, 0, 0, 0, 0, 0, 0, 0, 0, 0, 0, 0, 0, 0, 0, 0, 0, 0, 0, 0, 0, 0, 0, 0, 0, 0, 0, 0, 0, 0, 0, 0, 0, 0, 0, 0, 0, 0, 0, 0, 0, 0, 0, 0, 0, 0, 0, 0, 0, 0, 0, 0, 0, 0, 0, 0, 0, 0, 0, 0, 0, 0, 0, 0, 0, 0, 0, 0, 0, 0, 0, 0, 0, 0, 0, 0, 0, 0, 0, 0, 0, 0, 0, 0, 0, 0, 0, 0, 0, 0, 0, 0]
- **Raw Predicted Tokens:** [12, 18, 58, 1458, 7521, 11886, 2164, 7706, 876, 12090, 2, 2, 2, 2, 2, 2, 2, 2, 2, 2, 2, 2, 2, 2, 2, 2, 2, 7054, 2, 12, 2, 2, 2, 2, 2, 2, 2, 2, 2, 2, 2, 2, 2, 2, 2, 2, 2, 2, 2, 2, 2, 2, 2, 2, 2, 2, 2, 2, 2, 2, 2, 2, 2, 2, 2, 2, 2, 2, 2, 2, 2, 2, 2, 2, 2, 2, 2, 2, 2, 2, 2, 2, 2, 2, 2, 2, 2, 2, 2, 2, 2, 2, 2, 2, 2, 2, 2, 2, 2, 2, 2, 2, 2, 2, 2, 2, 2, 2, 2, 2, 2, 2, 2, 2, 2, 2, 2, 2]
- **Decoded Prediction Tree:** Cellular organisms, Archaea, Euryarchaeota, Methanomada group, Methanobacteria, Methanobacteriales, Methanobacteriaceae, Methanobrevibacter, Methanobrevibacter arboriphilus

### C.5 Example 5

- **Ground Truth Tree:** Cellular organisms, Eukaryota, Opisthokonta, Metazoa, Eumetazoa, Bilateria, Deuterostomia, Chordata, Craniata, Massilimicrobiota sp. An142, Gnathostomata (jawed vertebrates), Teleostomi, Euteleostomi, Actinopterygii, Actinopteri, Neopterygii, Teleostei, Osteoglossocephalai, Clupeocephala, Euteleosteomorpha, Protacanthopterygii, Salmoniformes (salmons and trouts), Salmonidae (salmonids), Salmoninae (trouts, salmons & chars), Oncorhynchus, Oncorhynchus tshawytscha (Chinook salmon) (Salmo tshawytscha)
- **Raw Ground Truth Tokens:** [12, 17, 53, 1418, 7435, 11764, 2030, 7481, 0, 8073, 0, 0, 0, 0, 5789, 0, 0, 0, 13743, 0, 0, 0, 8010, 0, 0, 0, 0, 0, 0, 0, 0, 0, 14711, 0, 5154, 3782, 0, 6167, 0, 0, 8328, 0, 0, 13891, 0, 0, 3237, 0, 0, 6317, 0, 0, 4786, 0, 0, 0, 0, 6275, 0, 0, 0, 4145, 0, 0, 5325, 0, 0, 8126, 0, 0, 0, 0, 0, 0, 0, 10584, 0, 0, 0, 0, 0, 3019, 0, 0, 0, 0, 0, 0, 2, 0, 0, 0, 0, 0, 0, 0, 0, 0, 0, 0, 0, 0, 0, 0, 0, 0, 0, 0, 0, 0, 0, 0, 0, 0, 0, 0, 0, 0]
- **Raw Predicted Tokens:** [12, 17, 53, 1418, 7435, 11764, 2030, 7481, 8073, 8073, 5789, 5789, 5789, 16, 5789, 16, 13743, 13743, 13743, 5789, 8010, 8010, 8010, 16, 14711, 14711, 14711, 12, 16, 12, 14711, 14711, 14711, 5154, 5154, 3782, 6167, 6167, 12, 8328, 8328, 13919, 13891, 13891, 5154, 6167, 3237, 4801, 6317, 6317, 4786, 4786, 4780, 14619, 6275, 6275, 6275, 6275, 12322, 4145, 4145, 4145, 13579, 5325, 5325, 14619, 8126, 8126, 12322, 10618, 10618, 3019, 10618, 8126, 7706, 10584, 3875, 3007, 3007, 3007, 3007, 3007, 2916, 2, 2, 2, 2, 2, 2, 2, 2, 2, 2, 2, 2, 2, 2, 2, 2, 2, 2, 2, 2, 2, 2, 2, 2, 2, 2, 2, 2, 2, 2, 2, 2, 2, 2, 2]
- **Decoded Prediction Tree:** Cellular organisms, Eukaryota, Opisthokonta, Metazoa, Eumetazoa, Bilateria, Deuterostomia, Chordata, Craniata, Massilimicrobiota sp. An142, Gnathostomata (jawed vertebrates), Teleostomi, Euteleostomi, Actinopterygii, Actinopteri, Neopterygii, Teleostei, Osteoglossocephalai, Clupeocephala, Euteleosteomorpha, Neoteleostei, Salmoniformes (salmons and trouts), Salmonidae (salmonids), Salmoninae (trouts, salmons & chars), Oncorhynchus, Oncorhynchus mykiss (Rainbow trout) (Salmo gairdneri)

## D Supplementary Figures

This appendix contains supplementary figures providing more detailed analysis of the model’s performance and its learned embedding space.

**Figure 6:**
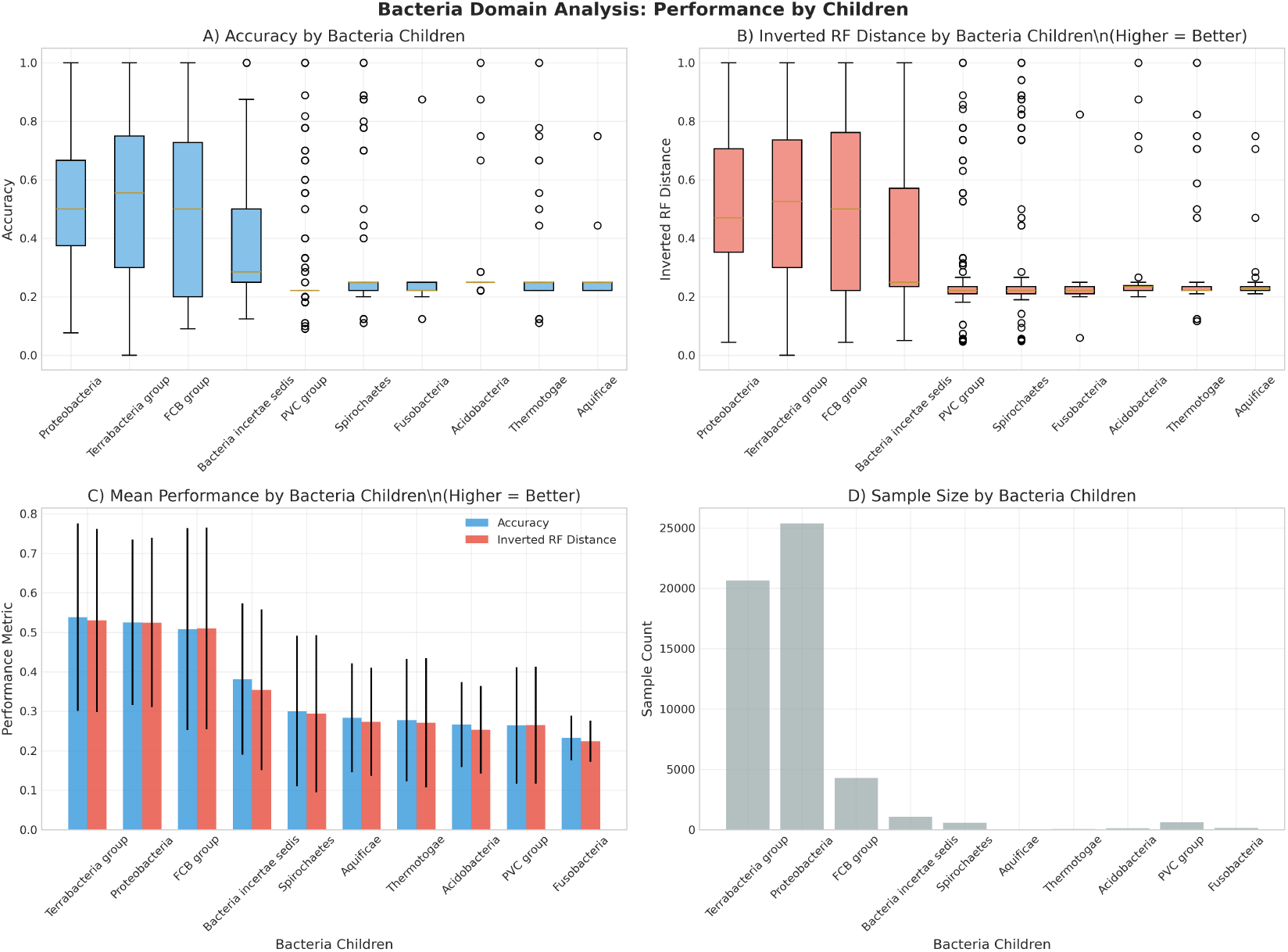
Performance Breakdown within Bacteria. This figure provides a granular analysis of performance for the major phyla (children) within the Bacteria domain. The four subplots show that performance metrics like Accuracy (A) and Inverted RF Distance (B) are strongly correlated with the sample size for each phylum (D). Well-represented phyla like Proteobacteria and Terrabacteria group show significantly better and more consistent performance than sparsely-sampled phyla.

**Figure 7:**
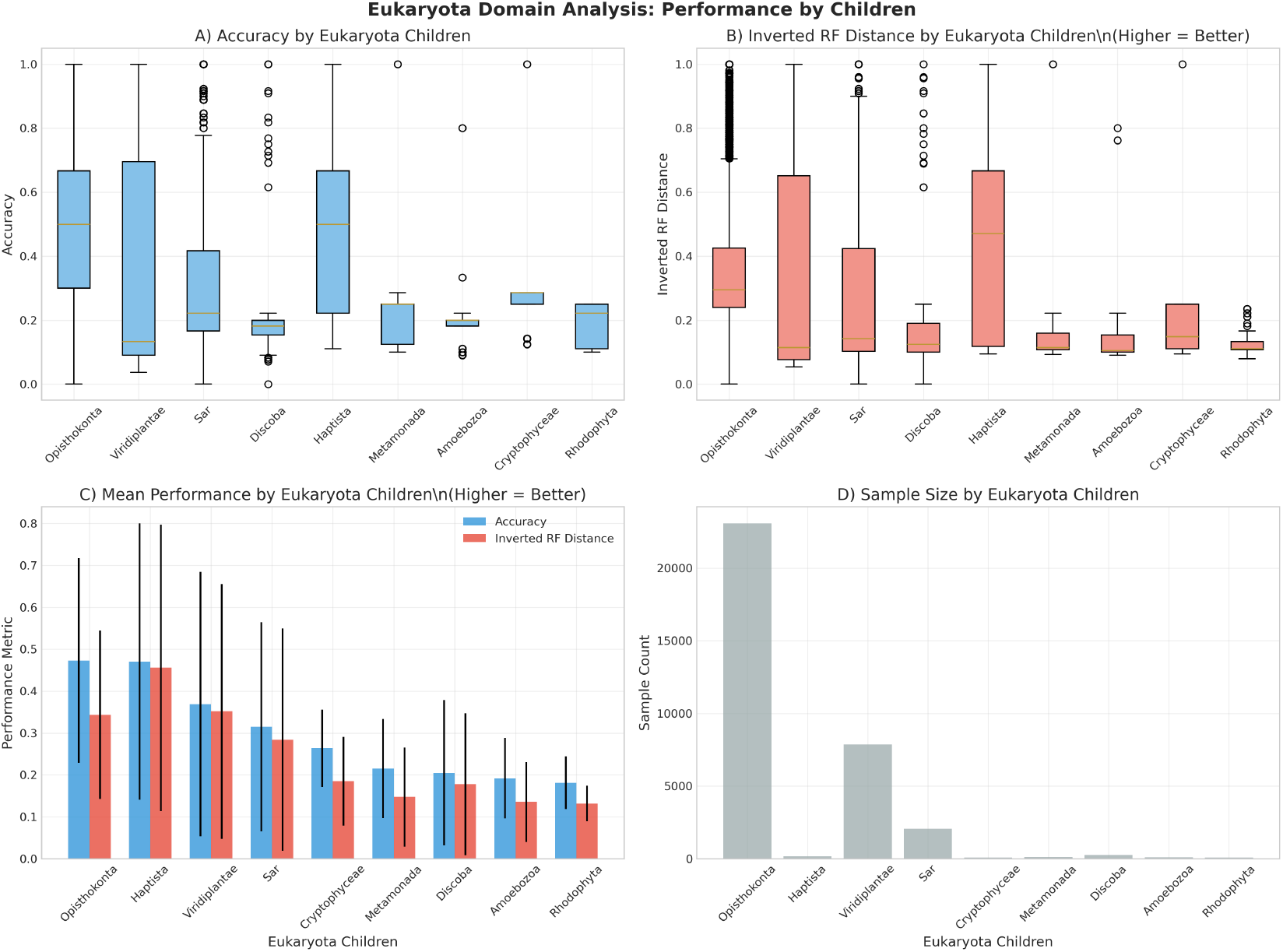
Performance Breakdown within Eukaryota. Similar to the analysis for Bacteria, this figure shows performance across major phyla within the Eukaryota domain. The dramatic difference in sample size (D), with Opisthokonta (which includes fungi and animals) being overwhelmingly dominant, directly translates to performance. Opisthokonta and Viridiplantae (plants) show the best performance, while other phyla with far fewer samples perform poorly.

**Figure 8:**
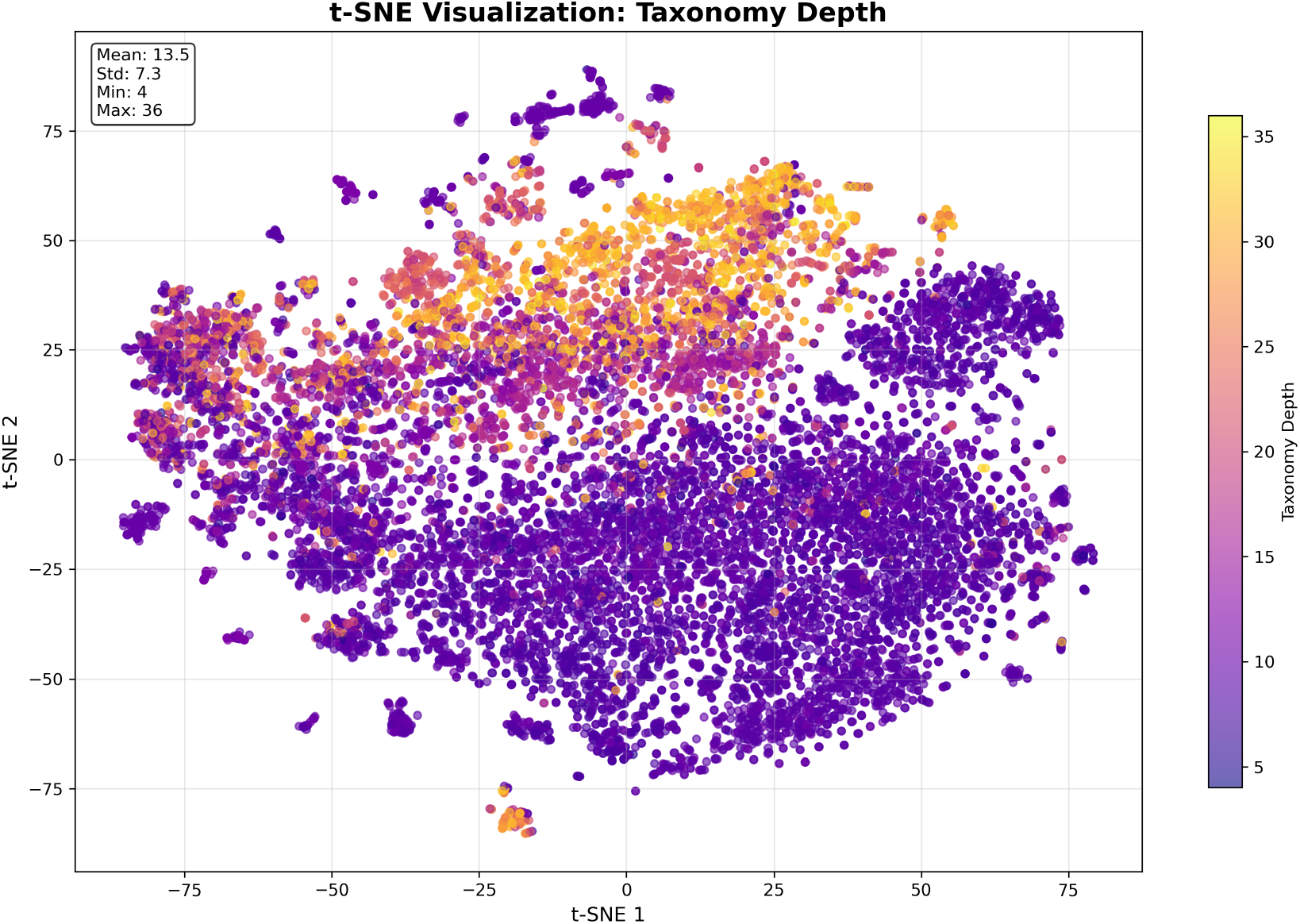
t-SNE of Embeddings by Taxonomy Depth. The learned protein embeddings show a clear global structure corresponding to the depth of the true taxonomic lineage, with shallower taxa (purple) and deeper taxa (yellow) occupying different regions of the space. This suggests the model learns a coarse representation of evolutionary depth.

**Figure 9:**
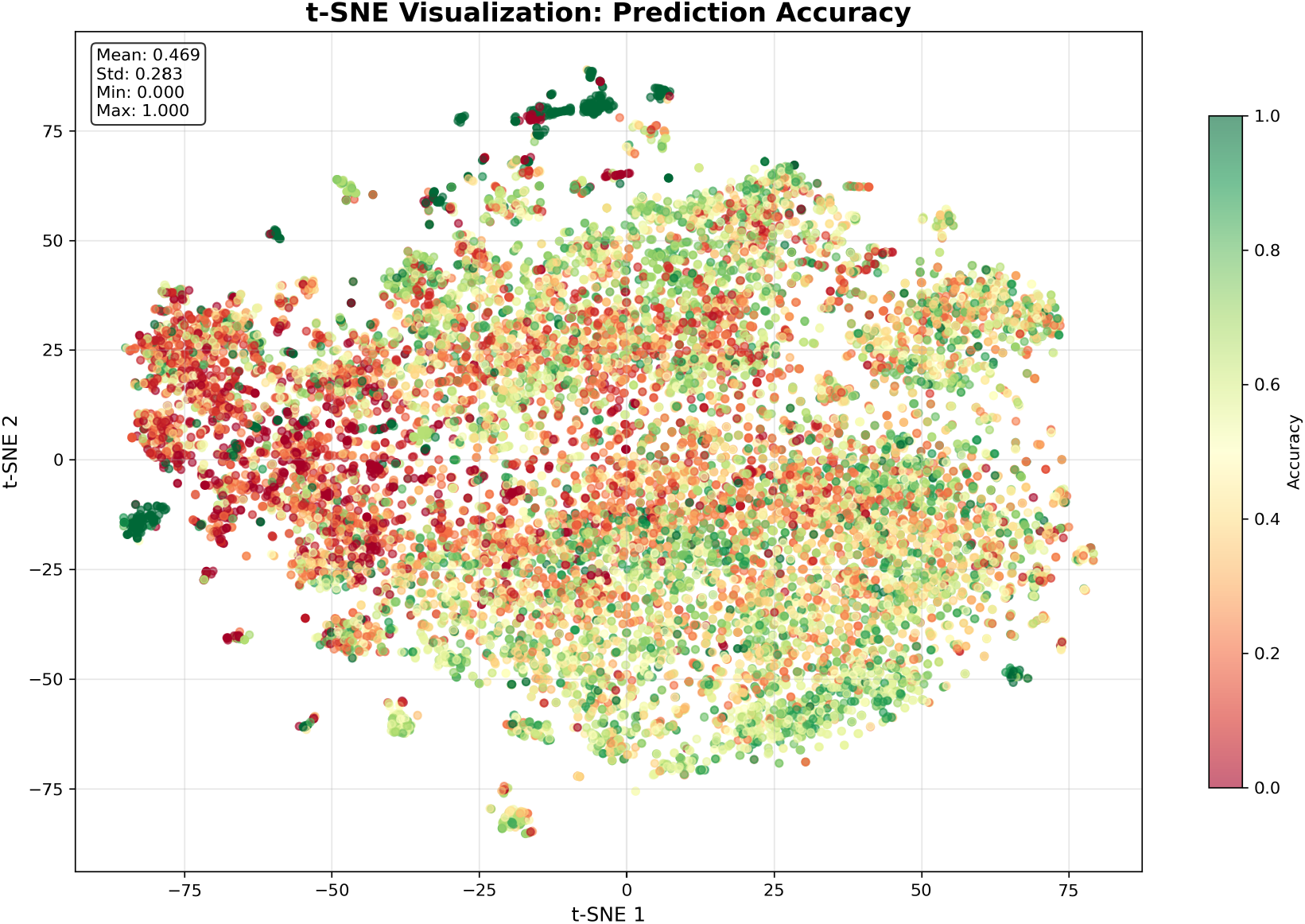
t-SNE of Embeddings Colored by RF Distance Error. This visualization highlights regions of the latent space where the model makes larger topological errors (yellow). These clusters may correspond to challenging or sparsely-sampled regions of the tree of life.

**Figure 10:**
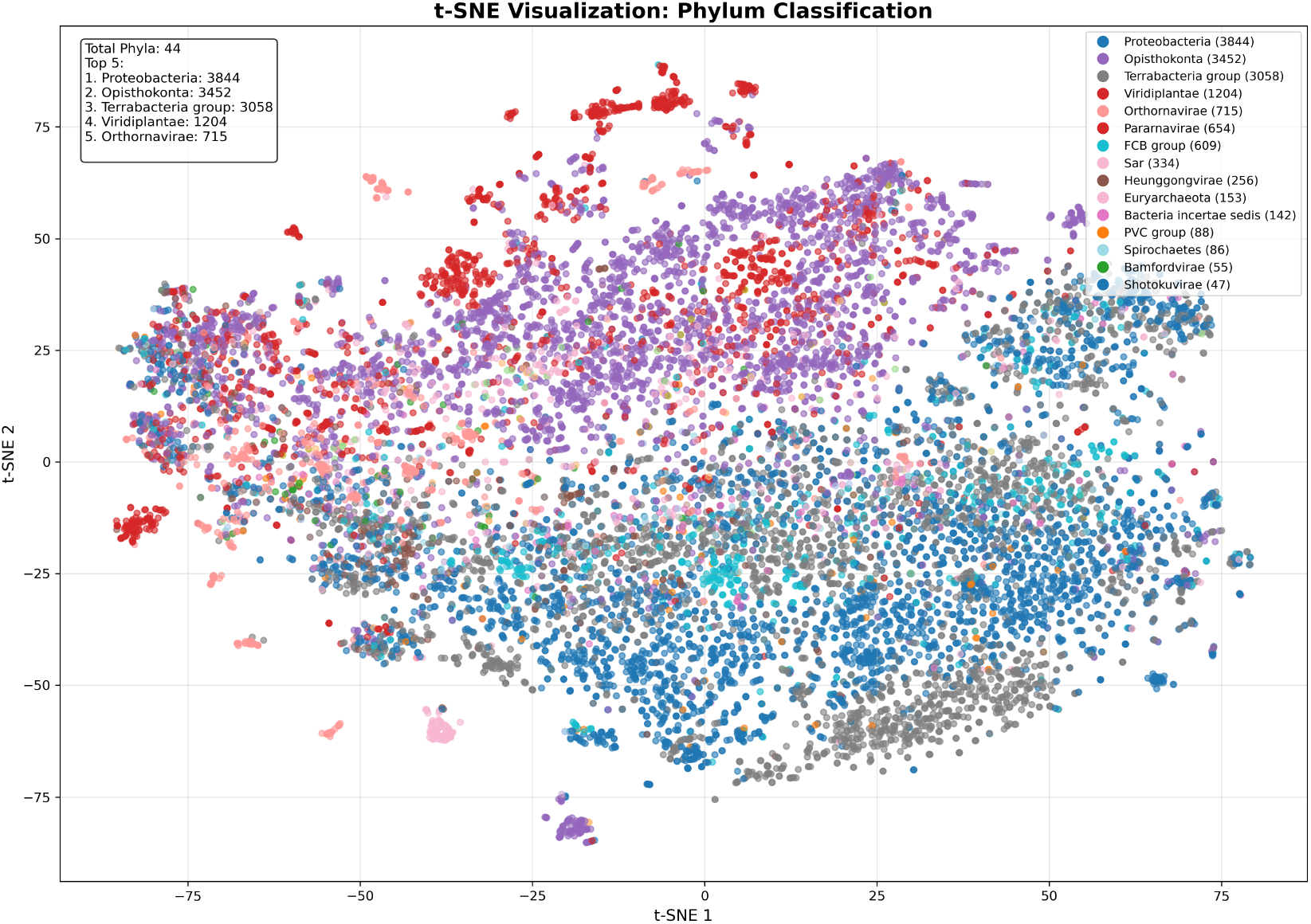
t-SNE of Embeddings by Phylum. This visualization shows the same protein embeddings from the evaluation set, but colored by their ground-truth phylum. The formation of distinct, large-scale clusters corresponding to major phyla (e.g., Proteobacteria, Opisthokonta) demonstrates that the model’s learned representation space is globally organized according to high-level taxonomic structure.

**Figure 11:**
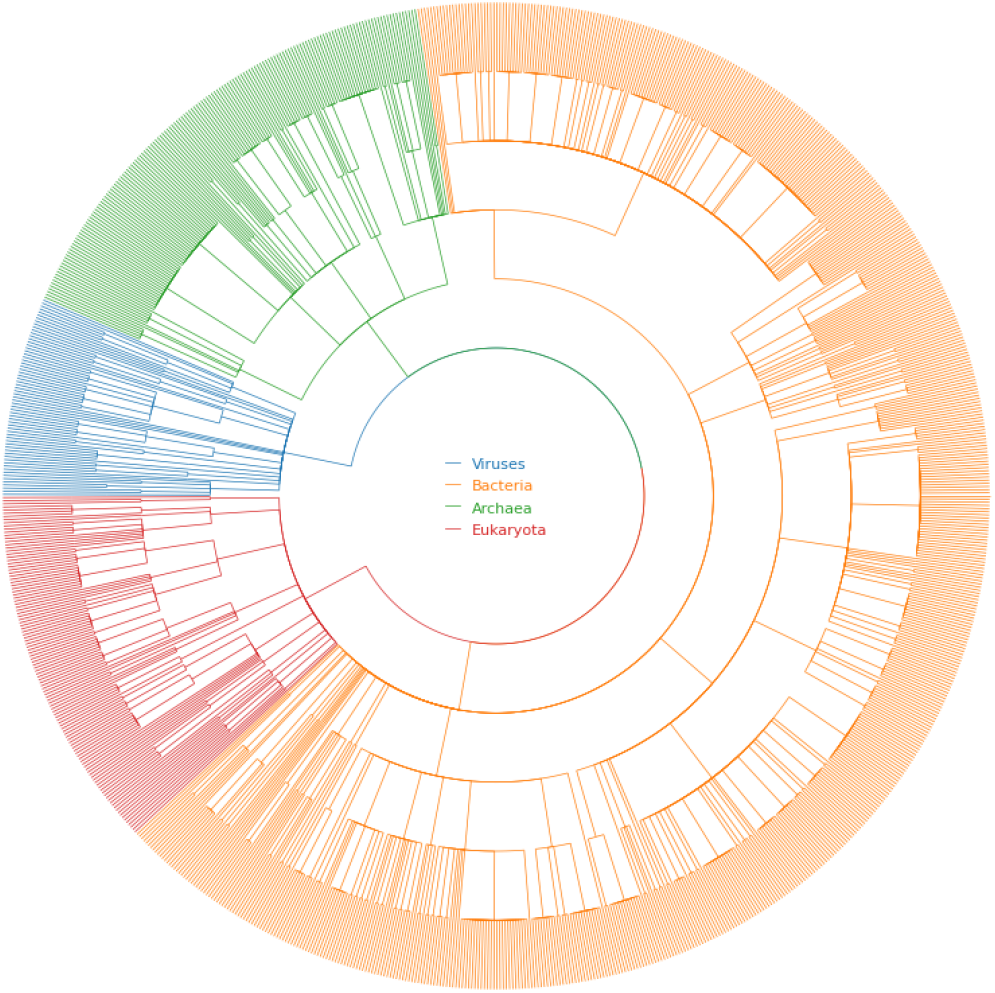
The Phylogenetic Tree of the Validation Set. This tree illustrates the immense diversity of the held-out test data, with branches colored by their top-level domain.

## Footnotes

2 Code available at https://github.com/msparsa/taxoformer

